# Heterogeneous flux capacity and oxygen sensitivity lead to subcellular ETC flux gradients in mouse oocytes

**DOI:** 10.64898/2026.06.29.735347

**Authors:** Maximilian Schwabe, Efe Ilker, Xingbo Yang

**Affiliations:** Cluster of Excellence Physics of Life (PoL), TU Dresden, Dresden 01062, Germany; Aix-Marseille Université INSERM, DyNaMo, Turing Centre for Living Systems, Marseille 13009, France

## Abstract

Mitochondria are metabolic hubs of the cell that provide energy and metabolites to meet the energetic, biosynthetic and signaling demands of the cell. Mitochondrial activities are characterized by the metabolic fluxes through their internal metabolic pathways. One of the most important mitochondrial metabolic pathways is the electron transport chain (ETC), where electron carriers such as NADH donate their electrons to oxygen to power mitochondrial respiration. Mitochondrial activities are dynamically and spatially regulated during organism development to ensure robust development. Recent work has revealed the existence of a subcellular ETC flux gradient within a single mouse oocyte, where mitochondria closer to the cell membrane display a higher ETC flux, but the mechanism underlying the formation of this gradient is unknown. In this work, we study the origin of the ETC flux gradients by modulating them through perturbations of external oxygen concentration and temperature. Interpreting the data with spatial kinetic modeling of mitochondrial respiration, we discover that the subcellular ETC flux gradient cannot be explained by reaction-diffusion of oxygen alone, but is a result of mitochondrial heterogeneity where mitochondria closer to the cell membrane display larger ETC flux capacity and lower oxygen sensitivity. Our work suggests that kinetically distinct subpopulations of mitochondria are spatially sorted according to their metabolic activities to form intracellular metabolic gradients.

## I. INTRODUCTION

Cells need energy to grow, proliferate and sustain life activities. Over a century of biochemistry research has revealed the details of metabolic pathways inside the cell [1]. However, most of our biochemistry knowledge is accumulated through population averages of cell metabolism or *in vitro* enzymology studies. These works are insufficient to reveal how metabolic activities are spatially regulated in living cells. Emerging evidence shows that the spatial patterning of metabolism plays a critical role in development by regulating cell migration and differentiation during embryogenesis [2–4]. To understand how cells form robust patterns in development, it is important to understand the mechanisms underlying the formation of spatial metabolic patterns and their functions.

Mitochondria are energy hubs of the cell and highly dynamic organelles. Their abundance, morphology, location and activities are actively controlled by the cell to meet energy and signaling demands. Mitochondria undergo fusion and fission, creating dynamic mitochondrial networks throughout the cell. Drp1-mediated mitochondria fission has been shown to facilitate follicle cell differentiation during *Drosophila* oogenesis [5]. Mitochondria are transported by molecular motors along actin filaments and microtubules [6]. Remodeling of actin filaments drives mitochondrial clustering that leads to up-regulated ATP production during mouse oocytes maturation [7]. Disrupting mitochondrial distribution in early mouse embryos compromises preimplantation development by inhibiting DNA replication and transcription [8].

Mounting evidence suggests that mitochondrial metabolism is spatially patterned and correlated with key developmental events [9, 10]. After the first cell differentiation event in mammalian embryo development, the trophectoderm cells display a higher oxygen consumption rate compared to the inner cell mass [11]. The spatial metabolic pattern exists even at the subcellular level: mitochondria closer to the meiotic spindle in mouse oocytes display a higher membrane potential than those further away, establishing a spatial gradient of mitochondrial metabolism within a single cell [12]. The mechanism underlying the formation of the metabolic gradients and their biological functions remain elusive. This knowledge gap is partly due to the lack of technique to measure mitochondrial metabolic fluxes with subcellular resolution.

Mitochondria provide energy in the form of ATP to sustain life activities. Nutrients and oxygen enter mitochondria and power the production of ATP through mitochondrial respiration. In short, nutrients such as pyruvate enter mitochondria and drive the production of high energy electron carriers NADH through the tricarboxylate acid (TCA) cycle. NADH donates electrons to oxygen through the mitochondrial electron transport chain (ETC). This process is coupled to proton pumping across the mitochondrial inner membrane, establishing the proton motive force (PMF) across the membrane. The PMF drives protons through a rotary molecule embedded in the mitochondrial inner membrane called ATP synthase to produce ATP. This process is termed Oxidative Phosphorylation (OXPHOS).

NADH is autofluorescent and hence is a natural biomarker of cell metabolism [13]. Fluorescence lifetime imaging (FLIM) of NADH enables the quantitative measurement of the concentrations of enzyme-bound NADH and free NADH by taking advantage of the fact that bound NADH displays a significantly longer fluorescence lifetime than free NADH. The changes of NADH fluorescence lifetimes and bound fractions are related to cancer progression [14], nutrient sensing [15] and embryogenesis [16–18]. Recently, a technique to measure fluxes through mitochondrial electron transport chain (ETC flux) in living cells with subcellular resolution has been developed [19]. This technique combines fluorescence lifetime imaging (FLIM) of NADH with biophysical modeling of NADH redox cycles to infer NADH oxidation flux in a label-free manner. Using this technique, a radially symmetric spatial gradient of ETC flux is discovered, where mitochondria closer to the plasma membrane display a higher ETC flux compared to those towards the center of the cell [19]. This observation raises the question: what are the key determinants of the spatial ETC flux gradients? Addressing this question could give insights into the mechanisms underlying the spatial regulation of metabolic activities, providing a critical step towards elucidating metabolic heterogeneities in development.

In this work, we study how the subcellular ETC flux gradients are established by perturbing external oxygen level and temperature, two major factors that impact the mitochondrial metabolic fluxes. The apparent Michaelis-Menten constant of mitochondria for oxygen is surprisingly low and well below 1 *µ*M for isolated mitochondria [20]. This is two orders of magnitude lower than the dissolved oxygen level of 210 *µ*M in air-saturated water at 37°C, suggesting a low oxygen sensitivity of mitochondria. However, it remains unclear if extreme hypoxia conditions could limit mitochondria respiration deep into tissues and cells, creating gradients in ETC flux as a result of oxygen reaction-diffusion. This requires quantitative measurement of oxygen gradients inside tissues and cells. More generally, if the ETC flux gradient is due to reaction-diffusion of oxygen, modulating the reaction rates of mitochondrial respiration could impact the decay length scale of the ETC flux gradients. Temperature is known to significantly impact respiration rate of organisms. An empirical relationship is the *Q*_10_ rule, which states that the respiration rate of an organism roughly doubles every 10 degrees of temperature increase [21]. Therefore, perturbing temperature provides a way to test if reaction-diffusion of oxygen provides possible explanation for the formation of ETC flux gradients.

In this paper, we investigate how the subcellular ETC flux gradients are established using mouse oocytes as the model system. We test if the reaction-diffusion of oxygen is responsible for the formation of the ETC flux gradients by measuring the response of the ETC flux gradients in response to perturbations of oxygen level and temperature. Interpreting the data with spatial kinetic modeling of mitochondrial respiration, we discover that reaction-diffusion of oxygen alone is not sufficient to explain the ETC flux gradients, and spatial heterogeneity in the mitochondrial flux capacity and oxygen sensitivity is required to explain the data. Our results suggest that the presence of spatial organization of kinetically distinct subpopulations of mitochondria within a single cell.

## II. RESULTS

### A. Inference of Subcellular ETC Flux Gradients from FLIM of NADH at Varying Oxygen Levels

Earlier work has revealed the existence of a subcellular ETC flux gradient within mouse oocytes arrested at Meiosis II (MII) ([19]). Since oxygen is the electron acceptor for the ETC (Figure 1A), here we first ask how external oxygen levels in the media (*c*_out_) impact the ETC flux gradient in the MII oocytes. We performed two-photon FLIM imaging of NADH across the equatorial plane of the oocytes and obtained the intensity image of NADH (Figure 1B left). Since NADH is enriched in mitochondria, we segmented the mitochondria from the cytoplasm using a pixel classification algorithm provided by ilastik based on the intensity contrast (Figure 1B right). We analyzed only the mitochondrial NADH throughout this paper. We performed FLIM of NADH across a range of external oxygen levels from 50 *µ*M to approximately 0 *µ*M by dropping oxygen level in the chamber hosting the oocyte from 5% to 0% partial pressure at 37°C (Methods and Materials). The oxygen drop was performed slowly enough to ensure equilibration between the oxygen level in the gas phase and the liquid phase that surrounds the oocytes. In our analysis, we divided the oocyte into 10 concentric regions and calculated the average concentrations of enzyme-bound NADH ([NADH_b_]) and free NADH ([NADH_f_]) as a function of distance to the oocyte center (Figure 1D-E, Methods and Materials). We used the NADH redox model to relate the free and bound concentrations of NADH to the ETC flux (*J*_ox_) via the below relation (Methods and Materials and Ref [19])

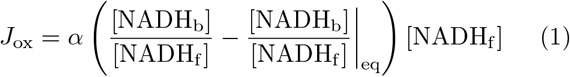

where “eq” denotes the value evaluated at equilibrium when *J*_ox_ = 0. *J*_ox_ has the unit of concentration per time, and characterizes the NADH oxidation rate per unit volume of mitochondria. In this work, the equilibrium condition is approximated at the lowest oxygen level. *α* is a temperature-dependent rate constant, whose values are calibrated by measuring oxygen consumption rate of the oocytes using respirometer (Methods and Materials). Applying Eq.(1) we obtain *J*_ox_ as a function of distance from the cell center at different external oxygen concentrations (Figure 1F). We observe that the *J*_ox_ displays a spatial gradient throughout all external oxygen levels, with a consistently higher level closer to the cell membrane and a lower level towards the cell center. The average *J*_ox_ level decreases when oxygen is dropped below a critical level.

**FIG. 1.**
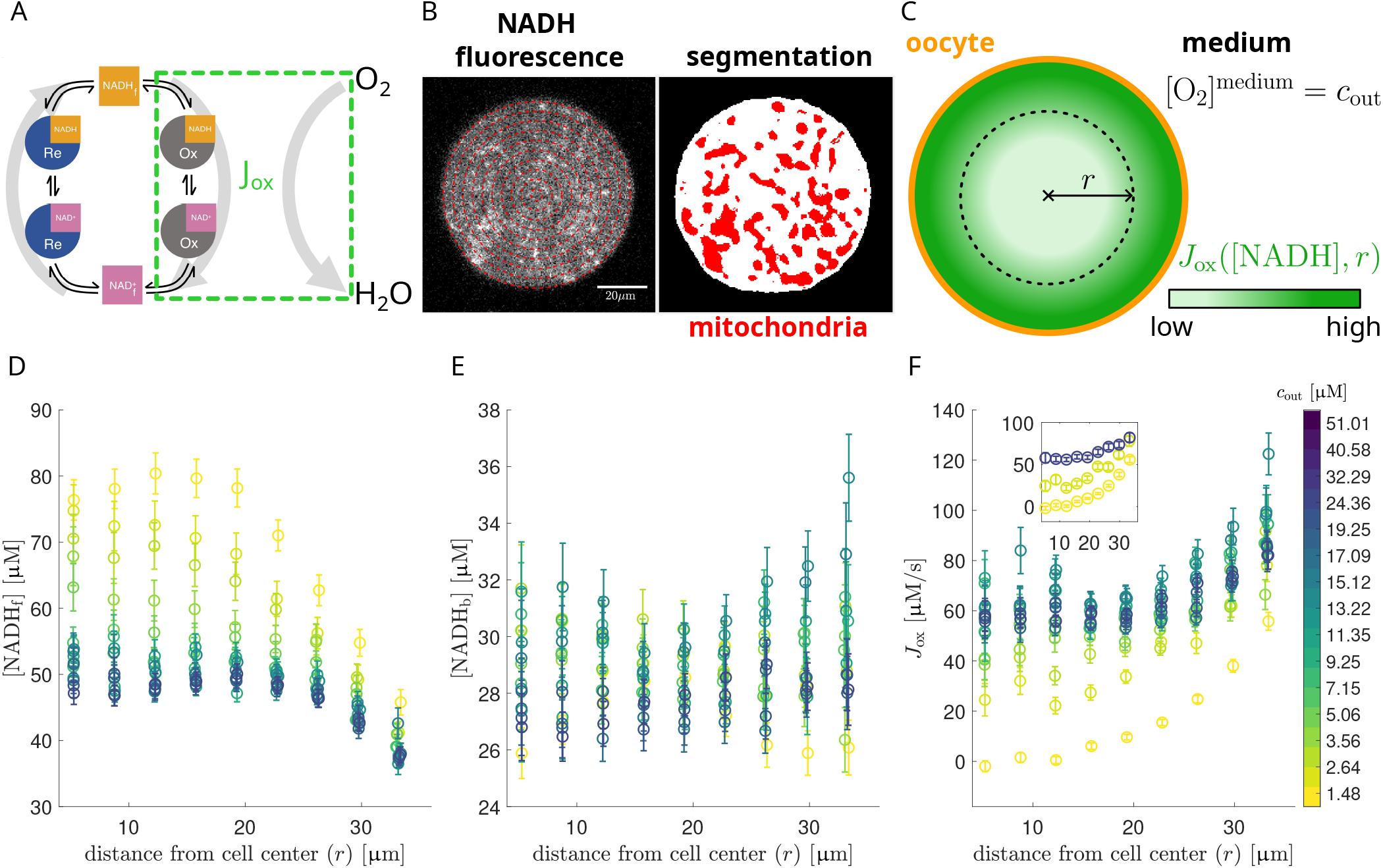
Gradients of subcellular ETC flux vary with external oxygen concentrations. **A:** Coarse-grained NADH redox model that relates the enzyme-bound and free NADH concentrations to the NADH oxidation flux, defined as the ETC flux *J*_ox_. The ETC flux is proportional to the mitochondrial oxygen consumption rate. The schematic was adjusted with permission from [19]. **B:** The autofluorescence intensity image of NADH inside an MII mouse oocyte measured by FLIM (left). In order to evaluate NADH fluorescence at subcellular resolution, the cell images are segmented into 10 concentric rings (indicated in red). Machine-learning based segmentation of the NADH autofluorescence signal into background and mitochondrial contributions (right). **C:** Schematic illustrating the spatial gradient of the NADH oxidative flux *J*_ox_ inferred from NADH concentrations at subcellular resolution. The midpoint radius of each segmented ring was used as the radial distance *r*. **D:** Concentration of free mitochondrial NADH as a function of distance from cell center. **E:** Concentration of bound mitochondrial NADH as a function of distance from cell center. **F:** Inferred oxidative flux *J*_ox_ as a function of distance from cell center. Different colored curves correspond to varying levels of external oxygen. The inset shows maximally different *J*_ox_ gradients at *c*_out_ = 0.49 µM, 1.48 µM and 51.01 µM. Results are computed by averaging over *N* = 67 individual oocytes. Error bars represent the corresponding standard error.

### B. Oxygen Gradients Alone Do Not Explain ETC Flux Gradients

Since oxygen diffuses from the boundary of the oocytes towards the interior of the cell, we ask if reaction-diffusion of oxygen explains the ETC flux gradient and its oxygen-dependence as observed in Figure 1F. Oxygen is the electron acceptor that receives the electron donated from NADH, therefore the ETC flux is proportional to oxygen consumption rate of the mitochondria (Figure 1A). First, we ask whether the ETC flux gradients can be captured by a first-order reaction kinetics of oxygen consumption. This would suggest that the ETC flux *J*_ox_(**r**) ∝*c*(**r**) where *c*(**r**) is the local oxygen concentration at a position **r** relative to the center of the oocyte.

To test this hypothesis, we first inferred the spatial distribution of oxygen concentration within the oocytes. In the cell interior, the oxygen is transported via diffusion and consumed in mitochondria following reaction-diffusion dynamics. Assuming a spherical symmetry of the problem we define the oxygen concentration *c*(**r**) = *c*(*r*) where *r* = | **r**|. At steady-state, the oxygen concentration *ĉ* should then satisfy

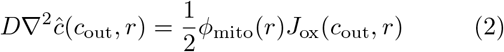

for 0 ≤ *r* ≤ *R* where *R* is the radius of the oocyte. Here, *ϕ*_mito_(*r*) is the local volume fraction of mitochondria and *J*_ox_(*c*_out_, *r*) is the local ETC flux depending on the external oxygen concentration *c*_out_. The factor 1/2 accounts for the fact that one O_2_ molecule is consumed per oxidation of two NADH molecules. Volume fraction of mitochondria *ϕ*_mito_(*r*) is measured by MitoTracker [19] and *J*_ox_(*c*_out_, *r*) is measured by FLIM of NADH as described in Section II A. Assuming spherical symmetry in Eq. 2 for *ĉ*(*c*_out_, *r*), we obtain

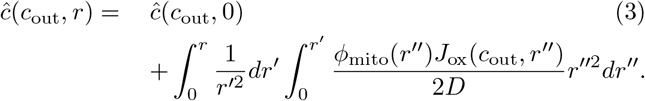

subject to boundary conditions

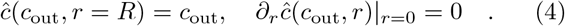

We solve *ĉ*(*c*_out_, *r*) through numerical integration of Eqs. (3) and (4). In Figure 2B, we plot the obtained intracellular oxygen levels relative to the external oxygen levels, i.e., *ĉ*(*c*_out_, *r*)*/c*_out_ as a function of distance from cell center for different external oxygen levels *c*_out_. This shows that the oxygen profiles are nearly flat at high *c*_out_ while steeper gradients form with decreasing *c*_out_. The integrated oxygen profiles are obtained with an oxygen diffusivity of D(*T* = 36°C) = 3320 *µ*m^2^*/*s as in water [22] and remain robust with respect to different choices of the oxygen diffusivity (See section I of Supplementary Materials). The magnitude of the oxygen gradients are similar to what has been calculated in bovine oocytes using a similar reaction-diffusion modeling approach [23].

**FIG. 2.**
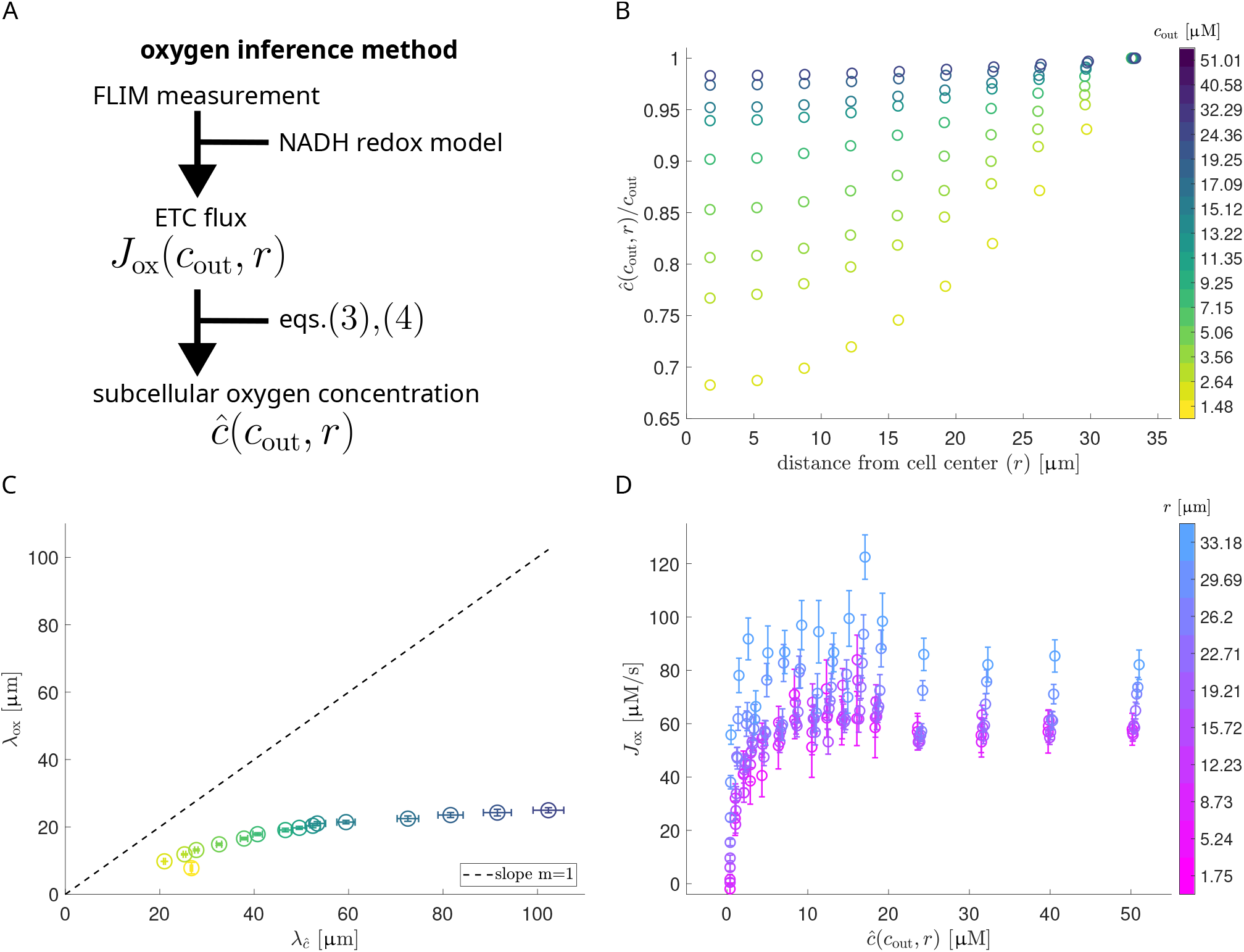
Reaction-diffusion of oxygen alone is not sufficient to explain the ETC flux gradients. **A:** Schematic of the pipeline used to estimate the subcellular oxygen concentration (*ĉ*) by numerically solving the steady-state reaction-diffusion equation of oxygen (eqn. 3, 4) with inferred spatial profile of the ETC flux (*J*_ox_). The inference of J_ox_ is based on the coarse-grained NADH redox model (Methods). Based on the assumption that mitochondrial consumption constitutes the main sink of oxygen inside the cell, spatial oxygen profiles can be obtained by numerical integration of eq.2 as in eqn.3. **B:** Inferred subcellular oxygen concentration as a function of distance from cell center. Different colors indicate varying external oxygen levels. Individual curves are normalized by corresponding oxygen level at the cell boundary. **C:** Fitting *n*(*r*) = (*λ/r*) ·sinh(*r/λ*) to normalized *J*_ox_(*c*_out_, *r*) and *ĉ*(*c*_out_, *r*) profiles at varying values of *c*_out_ yields characteristic decay lengths. The plot shows *J*_ox_ decay lengths *λ*_*ox*_ over *ĉ* decay lengths *λ*_*ĉ*_ corresponding to the same external oxygen level. Error bars correspond to the parameter standard error of the fits. There is a clear deviation from a linear relation with unit slope, i.e. *λ*_*ox*_ = *λ*_*ĉ*_ that would result from a reaction-diffusion system with linear oxygen consumption. **D:** Plotting oxidative flux *J*_ox_ as a function of inferred subcellular oxygen concentration *ĉ* shows a Michaelis-Menten like saturation behavior for high oxygen concentrations, which varies at different distances from cell center. An arbitrary nonlinear dependency of *J*_ox_ on oxygen concentration *ĉ* cannot explain the variability observed in the profiles at different distances. Error bars show the standard error of the oxidative flux across *N* = 67 oocytes considered.

To compare quantitatively ETC flux gradients (Figure 1F) and oxygen gradients (Figure 2B) at varying external oxygen levels *c*_out_, we determine their characteristic decay lengths. For an exponentially decaying profile along the radial direction *n*(*r*), a characteristic decay length *λ*_*n*_ can be obtained via the solution of 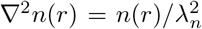. The Laplace operator ∇^2^ accounts for the spherical geometry of the problem and applies only in radial direction assuming spherical symmetry. The solution of this equation is a modified spherical Bessel function of the first kind, i.e., *n*(*r*)∝ (*λ/r*) sinh(*r/λ*)). Thus, by fitting the profiles normalized to their value at the cell boundary *r* = *R* to this function, we obtain the characteristic length scales *λ*_ox_ and *λ*_*ĉ*_. In Figure 2C we observe that *λ*_ox_ is not linear with respect to *λ*_*ĉ*_ and in fact saturates to an asymptotic value at high *c*_out_ while the oxygen profile flattens as indicated by a high value of *λ*_*ĉ*_. Even at low *c*_out_ levels, *λ*_ox_ ≠ *λ*_*ĉ*_ which rejects the first-order reaction kinetics hypothesis. In addition, the first-order reaction kinetics model cannot explain the ETC flux gradients with constant kinetic rates (see Section II of Supplementary Material).

We next test if the ETC flux is a unique non-linear function of the intracellular oxygen concentration following *J*_ox_ = *f* (*ĉ*) at any distance *r*. In Figure 2D, we plot *J*_ox_(*r*) within each concentric ring with an average distance of *r* to the oocyte center as a function of the intracellular oxygen concentration within that ring (*ĉ*(*r*)) throughout varying external oxygen levels *c*_out_. Each data set is color-coded according to the distance of its corresponding ring from the oocyte center. The lack of overlap among the data set indicates that *J*_ox_ is not a simple function of *ĉ* but instead exhibits additional spatial dependence.

Together, this data suggest that ETC flux gradients cannot be explained solely by intracellular oxygen levels and require an explicit spatial dependence such that *J*_ox_ = *f* (*ĉ, r*). Thus, we next turn our attention to space-dependent reaction kinetics.

### C. Spatially Heterogeneous Michaelis-Menten Model Captures the ETC Flux Profiles

The space-dependent *J*_ox_(*r, c*_out_)-*ĉ*(*r, c*_out_) relation indicates spatially heterogeneous reaction kinetics. Moreover, we observe saturation of *J*_ox_(*r, c*_out_) at high oxygen concentrations *ĉ*(*r, c*_out_) (Figure 2D). Simple reaction kinetics that can capture these aspect are described by a spatially heterogeneous Michaelis-Menten model which is given by:

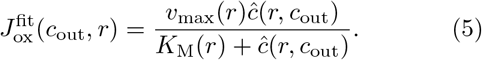

where *v*_max_(*r*) and *K*_M_(*r*) are respectively the space-dependent saturating flux value and the Michaelis-Menten constant (see section III of Supplementary Material for model derivation). We define the saturating flux *v*_max_ as the ETC flux capacity associated with oxygen saturation. The sensitivity of the ETC flux to the intracellular oxygen concentration is defined as the fractional change of 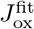 in response to fractional change of *ĉ*(*r, c*_out_)[24]:

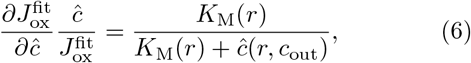

which increases with *K*_M_. We therefore identify *K*_M_ as a readout of oxygen sensitivity of the ETC flux.

We first fit *J*_ox_(*r*)-*ĉ*(*r*) curves at different distances *r* to the cell center as shown in Figure 3A. The *v*_max_(*r*) and *K*_M_(*r*) obtained from the fits display a spatial dependence where the flux capacity *v*_max_(*r*) is higher closer to the cell membrane and decreases towards the cell center, while the Michaelis-Menten constant *K*_M_(*r*) is lower closer to the cell membrane, indicating a lower oxygen sensitivity of the mitochondria, and increases towards the cell center (Figure 3C).

**FIG. 3.**
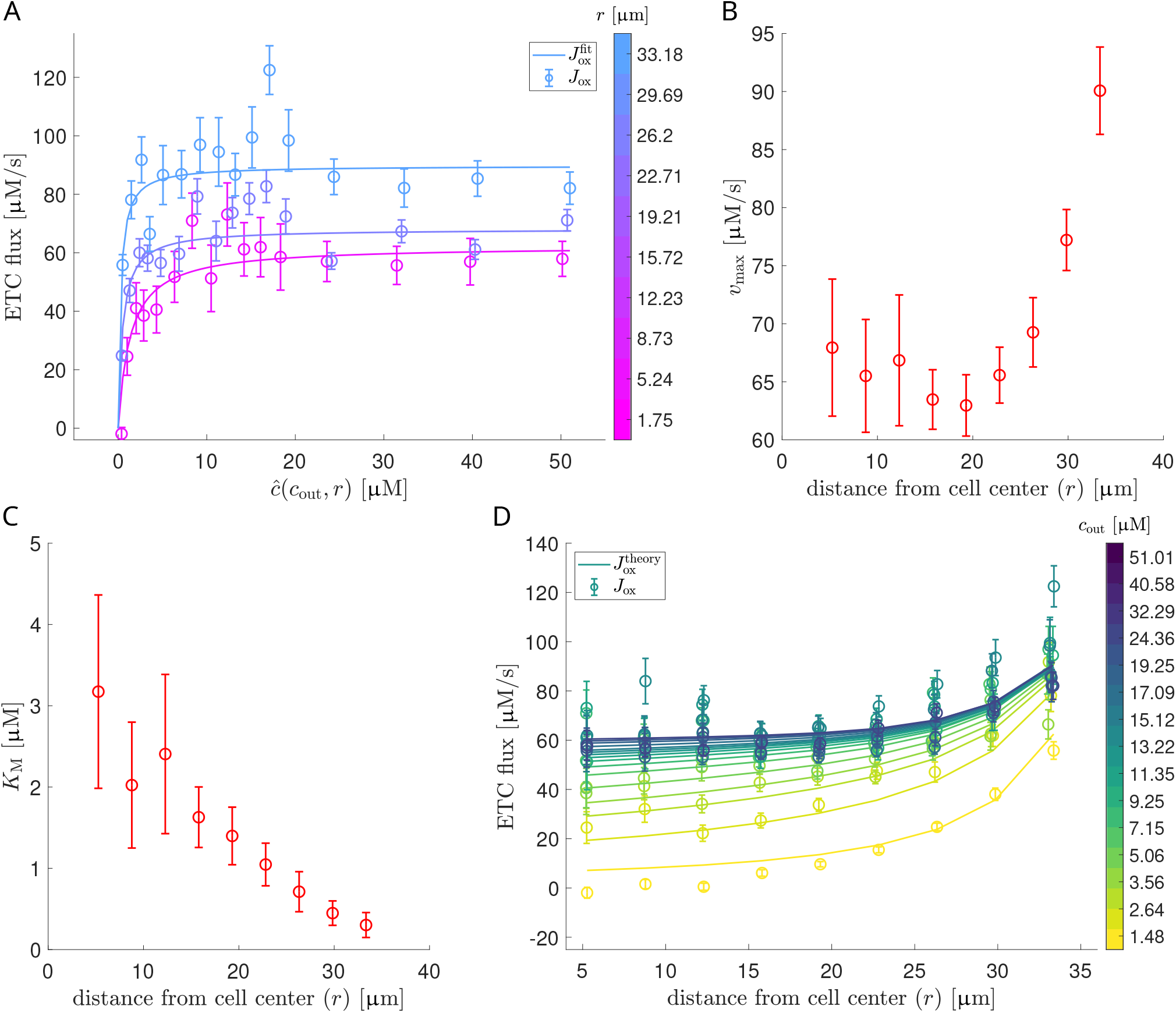
Reaction-diffusion with spatially-varying flux capacity and oxygen sensitivity explains the ETC flux gradients. **A:** Inferred (*J*_ox_) and fit 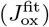 NADH oxidative flux (generally termed ETC flux) as a function of inferred subcellular oxygen concentrations. The oxygen dependency of *J*_ox_ is modeled as a Michaelis-Menten like relation with radially varying parameters (eqn 5). 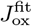 is computed based on fit parameter profiles *v*_max_(*r*), *K*_M_(*r*) and inferred oxygen *ĉ*(*c*_out_, *r*). Errorbars show the standard error across *N* = 67 oocytes. **B:** Spatial profile of the maximum ETC flux (*v*_max_) as inferred from Michaelis-Menten fits to *J*_ox_(*ĉ*(*c*_out_, *r*)) at varying *r*. Errorbars correspond to the parameter standard error of the Michaelis-Menten fit. **C:** Spatial profile of the apparent Michaelis-Menten constant (*K*_M_). **D:** Plot of ETC flux as a function of distance from cell center. Dots indicate empirical data (inferred *J*_ox_), solid lines the ETC flux predicted from oxygen reaction-diffusion with spatially varying parameters 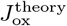, eqn.(8)). Errorbars show the standard error of *J*_ox_ across *N* = 67 oocytes.

We next apply the spatially heterogeneous Michaelis-Menten model in reaction-diffusion dynamics of oxygen for a consistency check. Inserting (5) in (2) yields:

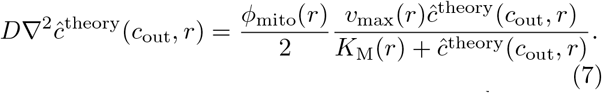

We solve this equation numerically for *ĉ*^theory^ using the fitted *v*_max_(*r*) and *K*_M_(*r*) obtained from *J*_ox_(*r*)-*ĉ*(*r*) curves (compare Figure 3B, C). For this, *v*_max_(*r*) and *K*_M_(*r*) are smoothed by approximate functional fits (exponential for *v*_max_(*r*), linear for *K*_M_(*r*)). From the oxygen profiles calculated this way, we predict the ETC flux as a function of distance to cell center using equation (5) at different external oxygen concentrations.

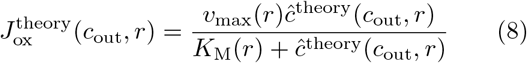

The predicted ETC flux gradients agree with the experimental measurements (Figure 3D). This suggests that oxygen reaction-diffusion with consumption kinetics characterized by a spatially-varying flux capacity and oxygen sensitivity explains the ETC flux gradients.

### D. Capacity and Oxygen Sensitivity of ETC Flux Increase with Temperature

Although the total metabolic rate of an organism is known to be sensitive to temperature variations [25], it is unexplored how metabolic flux gradients depend on temperature. Our experimental system can provide insights to this question as we can vary the temperature of the system and make a comparative analysis. In mouse oocytes, if the ETC flux gradients are driven by reaction-diffusion of metabolites, such as oxygen, that feed into mitochondrial respiration, we expect the decay length of the ETC flux gradients to be sensitive to temperature variations when the metabolic rate and diffusivity of the metabolite display different temperature-dependence. Therefore, temperature variation experiments provide another route to test the mechanism underlying the formation of spatial metabolic gradients.

We first study the impact of temperature on the average *J*_ox_ of the oocytes. MII oocytes are incubated at T=22°C, 28°C, 31°C and 36°C. ETC flux at each temperature are predicted using Equation 1, where the calibration factor *α*(*T*) is adjusted for different temperatures (Materials and Methods). Then, using the free and bound NADH concentrations at different temperatures (see section IV of Supplement Material for profiles of free and bound NADH), we predict ETC flux with absolute units across all the temperatures and averaged over three different external oxygen concentrations. The averaged *J*_*ox*_ (Figure 4A) increases with temperature across all three external oxygen concentrations from 20 *µ*M to 50 *µ*M, which corresponds to the saturated regime of the *J*_ox_. Next, we explore the impact of temperature on the

**FIG. 4.**
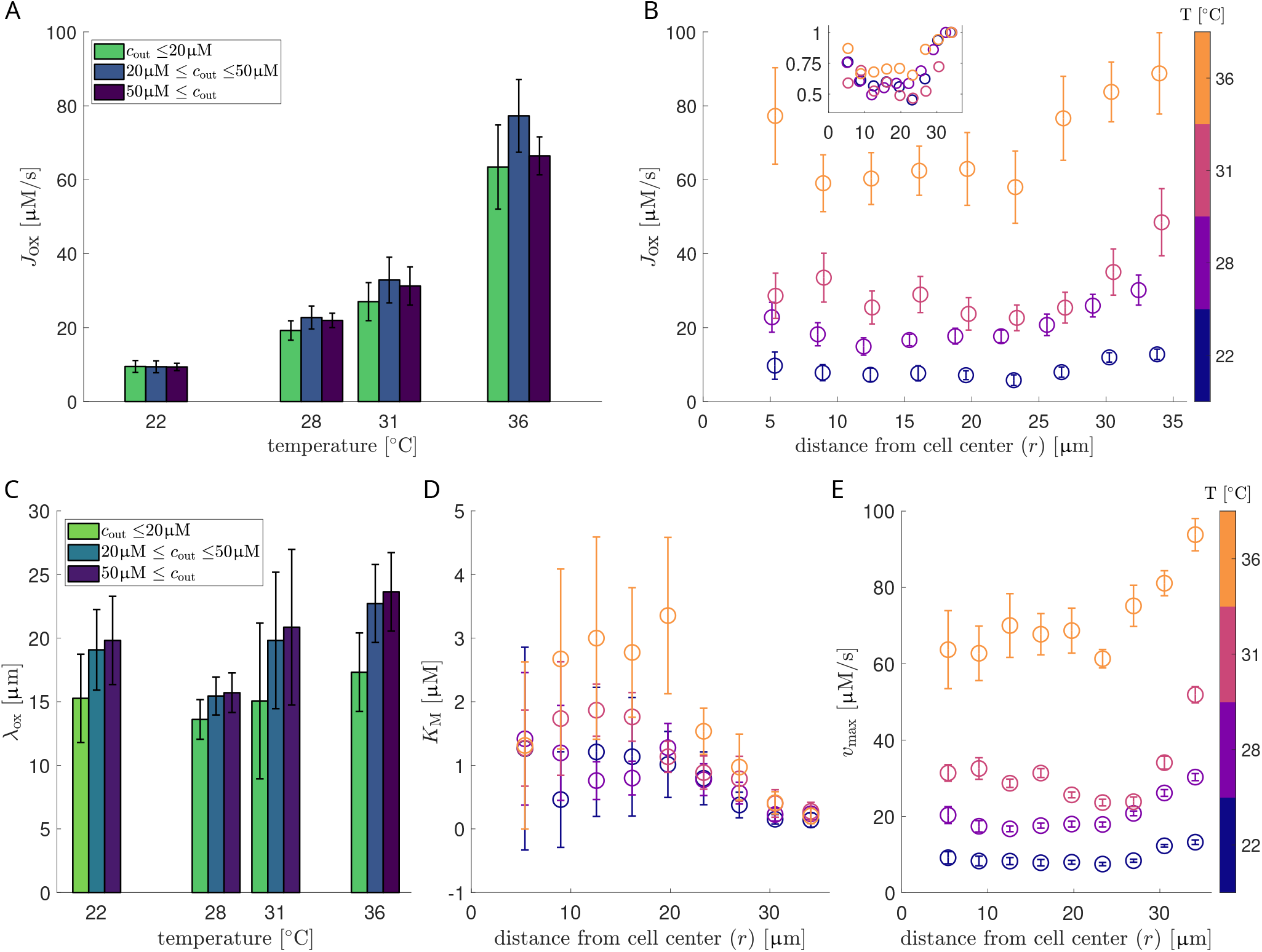
Temperature increases ETC flux capacity and oxygen sensitivity but does not impact decay length scale of flux gradients. **A:** Whole-cell averaged *J*_ox_ at different temperatures. Colors indicate averages over values from different regimes of external oxygen concentrations. For all external oxygen concentrations, the cell-averaged *J*_ox_ increases significantly with temperature. Errorbars correspond to the standard error of *J*_ox_ over oocytes considered at different temperatures. **B:** Inferred oxidative flux *J*_ox_ as a function of distance from cell center at four different temperatures averaged over external oxygen concentrations from a range of 20 *< c*_out_ *<* 50*µ*M. The inset shows the *J*_ox_ at different temperatures normalized by the respective flux at the boundary. Errorbars show the standard error of *J*_ox_ across different numbers of oocytes (*N*) for each temperature (*T* = 22°C: *N* = 10, *T* = 28°C: *N* = 13, *T* = 31°C: *N* = 10, *T* = 36°C: *N* = 8) **C:** Decay lengths of *J*_ox_ gradients estimated by fitting *n*(*r*) = (*λ/r*) ·; sinh(*r/λ*) to normalized *J*_ox_(*c*_out_, *r*) profiles at varying values of *c*_out_ and different temperatures. Colors indicate decay lengths obtained at different external oxygen concentrations. *J*_ox_ gradient decay lengths are insensitive to temperature. Errorbars show the average uncertainty of the exponential fits used to extract decay lengths. **D, E:** Spatial profiles of oxygen sensitivity (*K*_M_) and ETC flux capacity (*v*_max_) at different temperatures. Both *v*_max_ and *K*_M_ increase with temperature. Errorbars correspond to the parameter standard error of the Michaelis-Menten fit.

ETC flux gradients in mouse oocytes. In Fig.4B, we observe that the ETC flux gradients persist across all temperatures. To quantify the impact of temperature on the ETC flux gradients, we fitted the decay length scales of the ETC flux gradients at different temperatures and the same three external oxygen concentrations. Remarkably, the decay length scales do not change significantly with temperature (Figure 4C). We also observe that at the highest oxygen level and from 22°C to 36°C, the oxygen diffusivity increases by a factor of 1.5 [22] while the average ETC flux increases by a factor of 7.1 (Figure 4A). Reaction-diffusion length would scale proportionally to 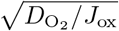 at similar oxygen levels which would predict that ETC flux decay length at 22°C is larger than the one at 36°C by a factor of 2.2. However, we observe in experiments that at the highest oxygen level the ETC flux decay length at 22°C is 19.8 ± 3.5 *µ*m whereas at 36°C it is 23.6 ± 3.1 *µ*m, displaying no significant difference (Figure 4C). The low sensitivity of the ETC flux decay length to temperature further supports that simple reaction-diffusion kinetics of oxygen is not sufficient to explain the ETC flux gradients.

Finally, we test the temperature dependence of the spatially-dependent kinetic parameters *v*_max_(*r*) and *K*_M_(*r*). *K*_M_ shows persistent spatial gradients across all temperatures, and the value increases with temperature especially at the center, which means that the oxygen sensitivity of the mitochondria increases with temperature especially at the oocyte center (Figure 4D). The steepness of the gradient of *K*_M_ increases with temperature, suggesting that mitochondria at different locations have different temperature sensitivities. The flux capacity *v*_max_(*r*) displays spatial gradients resembling those of the ETC flux, with an increasing average value with temperature (Figure 4E), suggesting that the ETC flux gradient at saturating oxygen level is due to a spatially varying flux capacity.

## III. CONCLUSIONS AND DISCUSSION

In this work, we aimed to gain insights into spatiotemporal patterning of metabolic activity in cells which is integral to embryo development [2–4]. For this, we use mouse oocyte as a model system for which recent work has revealed the existence of a subcelllular ETC flux gradient [19]. We systematically tested the impact of oxygen levels and temperature on the ETC flux gradient with the aim of elucidating the mechanisms underlying the formation of this flux gradient.

Our analysis combining theory and experiment demonstrates that the ETC flux gradient cannot be explained by the intracellular oxygen gradients as a result of simple reaction-diffusion of oxygen. Instead we observe that the ETC flux is not a unique function of intracellular oxygen levels, but displays additional spatial dependency that indicates spatially heterogeneous reaction kinetics. To capture this behavior, we propose a Michaelis-Menten model with spatially-varying flux ca-pacity and oxygen sensitivity. This model quantitatively explains the ETC flux gradients as a result of the higher ETC flux capacity and lower oxygen sensitivity of mitochondria localized closer to the cell membrane. These observations suggest the possible existence of heterogeneous subpopulations of mitochondria within a single cell. Latest research shows that heterogeneous subpopulations of mitochondria indeed exist within a single cell, where some mitochondria are specialized in ATP production while others focus on biosynthesis [26]. Our previous work [19] demonstrates that the ETC flux gradient persists even after the inhibition of ATP synthase of mitochondria, suggesting that the mitochondrial heterogeneity lies outside of ATP synthesis.

One hypothesis for the molecular mechanism responsible for the ETC flux gradient could be a heterogeneous distribution of mitochondrial enzyme concentrations or a heterogeneous enzyme catalysis rate. In section III of Supplementary Material we show a derivation of the spatial kinetic model based on Michaelis-Menten kinetics with spatially heterogeneous enzyme concentrations and catalysis rates. This picture would suggest that mitochondria closer to the cell membrane could possess a higher level of ETC complexes or uncoupling proteins compared to those in the cell interior. Antibody staining targeting specific ETC complexes or spatial proteomic quantification are needed to test this hypothesis. Spatially-dependent association with other cell organelles could contribute to the heterogeneity in enzyme catalysis rate. Mitochondria associated with lipid droplets display different bioenergetic profiles compared to isolated mitochondria [27]. In oocytes, mitochondria close to the meiotic spindle display higher mitochondrial membrane potential [12]. In MII oocytes as studied in this work, the meiotic spindle is located asymmetrically near the cell membrane [28]. Since the equatorial plane is randomly chosen across many oocytes, the potential angular dependence of mitochondrial metabolic state associated with the position of the spindle is expected to average out, leaving a radially-symmetric metabolic gradient as observed in this study. To study how other cellular structures impact the mitochondrial metabolic gradients requires a combination of confocal imaging of these structures together with ETC flux mapping. Another possible mechanism for the establishment of the ETC flux gradient is from reaction-diffusion of signaling molecules, such as calcium and reactive oxygen species (ROS), both of which are known to modulate mitochondrial metabolism[29, 30]. Biosensors capable of measuring calcium and ROS spatial gradients are required to test this hypothesis.

Moreover, our work reveals a spatially heterogeneous mitochondrial oxygen sensitivity, related to the apparent Michaelis-Menten constant *K*_M_. Remarkably, *K*_M_ takes a very small value, on average about 1*µ*M, indicating that the half maximum ETC flux occurs around 1*µ*M of oxygen concentration, which is extremely low compared to the oxygen concentration in air-saturated aqueous solution at 37°C, which is around 210*µ*M. This suggests that mitochondria have a very low oxygen sensitivity, consistent with early studies on isolated mitochondria [20]. It remains a long-standing question as to what factors account for this apparent low oxygen sensitivity of mitochondria. The known enzyme oxygen binds to in the electron transport chain of mitochondria is Cytochrome c Oxidase, also known as the Complex IV [1]. *In vitro* assay of purified Cytochrome c Oxidase shows that the apparent dissociation constant of oxygen to Cytochrome c Oxidase is 280*µ*M [31], far greater than the value in isolated mitochondria or intact cells. This discrepancy is likely an emergent property arising from the kinetics and thermodynamics of the entire ETC, and not attributable to a single enzymatic step. Existing theories to explain the low oxygen sensitivity include kinetic trapping [32] and co-substrate compensation [33]. However, these theories do not explain the observations that the apparent *K*_M_ changes with the energetic state of the mitochondria. Particularly, *K*_M_ has been shown to decrease in uncoupled mitochondria where proton leakage dominates [34]. Consistent with this earlier study, in our previous work [19], we have shown that mitochondria closer to the cell membrane have a lower mitochondrial membrane potential and a higher proton leak flux, which correlates with the lower *K*_M_ observed in this study. This could suggest that mitochondria close to the cell membrane is less efficient in terms of ATP production compared to those at the cell center. To quantitatively understand the correlation between *K*_M_ and energetic state of the mitochondria in a spatially-dependent manner is a new question motivated by our research.

Temperature is an important controller of metabolic rate. An empirical rule determines that metabolic rate doubles approximates every 10 degrees increase of temperature, termed the Q10 rule [21]. This increase of metabolic rate is correlated with the increase of developmental tempo in organisms such as *Drosophila* [35]. How temperatures regulate developmental rate through metabolic rate is an open question. Our work shows that the average ETC flux of the oocytes increases by a factor of 7.1 from 22°C to 36°C, which is more than what is predicted by the Q10 rule. In contrast, the decay length of the ETC flux gradient does not change significantly with temperature. This result suggests that the rates of mitochondrial respiration respond uniformly to temperature. However, the *K*_M_ gradient becomes sharper with increasing temperature, demonstrating that the oxygen sensitivity responds in a spatially-dependent manner to temperature. These observations suggest that the ETC flux capacity and mitochondrial oxygen sensitivity are potentially controlled in a thermodynamically distinct manner in living cells. More detailed characterization of the ETC in combination with thermodynamic modeling is required to reveal the mechanism.

We note that while our work provides the first systematic study of the origin of a subcellular metabolic flux gradient and opens up various new questions, it has its own limitations. The intracellular oxygen levels are inferred from the reaction-diffusion equation of oxygen, with a oxygen diffusivity of D(*T* = 36°C) = 3320 *µ*m^2^*/*s as in water [22]. Since measurement of oxygen diffusivity in living cells is lacking, the inferred intracellular oxygen gradients could be subject to improvement. However, our results remain robust to variations in oxygen diffusivity (see section I of Supplement Material). Direct approaches to measure intracellular oxygen levels are generally lacking. Despite of the existence of oxygen-sensitive dyes or FRET-based biosensors [36, 37], calibrating these sensors quantitatively and excluding confounding factors to achieve the enough sensitivity to resolve subcellular oxygen gradient remains to be a challenge. Our measurement so far, due to limited imaging speed of FLIM of NADH, is constrained to 2D. To reveal the full 3D pattern of the ETC flux, 3D FLIM of NADH needs to be implemented. In mouse oocytes, mitochondria are known to accumulate near meiotic spindle and display distinct metabolic state [12, 28]. Therefore, the spindle could be a contributor to the ETC flux gradients.

Overall, our work reveals that the subcellular ETC flux gradients in mouse oocytes can be explained by a spatially heterogeneous flux capacity and oxygen sensitivity. This suggests a spatial organization of mitochondria populations sorted according to their biochemical activities. The functions of these subcellular flux gradients remain to be explored. Our framework combining a subcellular flux inference technique with quantitative metabolic perturbations could help answering questions regarding the mechanisms and functions of these ETC flux gradients in development.

## IV. MATERIALS AND METHODS

### Culturing of mouse oocytes

Frozen MII mouse oocytes (Strain B6C3F1) from EmbryoTech were used for this work. Oocytes were thawed and cultured in AK-SOM media (MilliporeSigma) covered with mineral oil (VitroLife) and incubated at 37C with 5% CO_2_ level and air saturated oxygen level. For imaging, oocytes were transfered to a 2uL AKSOM droplet in a 35mm glass bottom dish (FluoroDish, WPI) covered with 400-500uL Mineral oil to minimize gas exchange time for the subsequent oxygen drop experiment. The glass bottom dish was placed in a sealed Ibidi chamber during imaging with temperature maintained at 37C and CO_2_ maintained at 5%. Oxygen level was maintained at 5% in the chamber before oxygen drop.

### Two-photon FLIM of NADH measurements

FLIM data were collected with a home-built two-photon scanning confocal microscope. The system is with a 40 × 1.25NA water immersion Nikon objective, Becker and Hickle Time Correlated Single Photon Counting (TCSPC) acquisition system with HPM-100-40 GaAsP hybrid PMT detector and a 80 MHz pulsed MaiTai DeepSee Ti:Sapphire laser (Spectra-Physics). NADH autofluorescence was excited at 750 nm wavelength with a power of 3 mW measured at the objective with parked laser beam. The scanning area is 512 × 512 pixel with a pixel size of 420 nm and a pixel dwell time of 4.09 us. The NADH autofluorescence was collected with a 460/50nm filter. NADH fluorescence decay curves were collected at each pixel.

### Oxygen drop experiment

After 15 mins of equilibration of the oocytes at 5% O_2_ and 5% CO_2_ in the Ibidi chamber, oxygen level in the chamber were continuously dropped from 5% to approximately 0% by manually mixing two pressurized gas tanks (Airgas) with 5% O_2_, 5% CO_2_ balanced with nitrogen and 0% O_2_, 5% CO_2_ balanced with nitrogen. Gas from both tanks were mixed through two tubes connected with a y-splitter and directed into the sealed Ibidi chamber. The oxygen level in the chamber was monitored in real time by directing the tube coming from the outlet of the chamber into a gas sensor (LOX-O2, Gaslab) and recorded via the Gaslab software. Given the existence of a finite offset of the sensor reading, the recorded oxygen level was corrected by subtracting the value recorded when directly pumping in gas from the 0%O_2_ gas tank into the sensor. The oxygen level in the chamber was continuously dropped from 5% to close to 0% as measured by the offset-subtracted sensor over a course of 30 mins, which is much longer than the diffusion time scale of oxygen through the thin layer of oil covering the media droplet housing the oocytes (∼5min as calibrated by oxygen sensitive Ruthenium dye when oxygen is depleted almost instantaneously in the gas phase). Hence the oxygen level can be considered in quasi-equilibrium between the gas phase and the liquid phase. Applying Henry’s law, we predicted a corresponding drop from 50 *µ*M to 0 *µ*M of dissolved oxygen concentration in the media at 37°C and 1 atm from the oxygen partial pressure in the chamber (at different temperature, the solubility is adjusted accordingly). FLIM of NADH was performed during the entire oxygen drop and the subsequent recovery of oxygen back to 5%. The viability of the oocytes were confirmed by the fact that the FLIM parameters returned to the pre-drop level after oxygen recovery.

### Temperature sweep experiment

Temperature at the droplet housing the oocytes was controlled by tuning the temperature of the Ibidi lid and plate together with an objective heater. The temperature at the droplet was calibrated with a thermocouple at 22°C, 28°C, 31°C and 36°C by adjusting the temperature control settings of the Ibidi chamber and the objective heater. The same settings were then used to maintain the temperature at 22°C, 28°C, 31°C and 36°C for the droplets housing the oocytes. FLIM of NADH was performed at each temperature for about 1 hour including oxygen drop and recovery after the temperature stabilizes. The oocytes survived the entire experiment and the change of NADH FLIM parameters during oxygen drop is reversed upon oxygen recovery, suggesting the reversibility of the experiment. Oxygen drop experiment was performed at each temperature following the same protocol as described above but with correction of oxygen solubility *µ*_ox_(*T*) at different temperatures (compare [22]). The experimental setup measures oxygen partial pressure *p* in the medium relative to 21% of oxygen in the gas phase. Following Henry’s law, one arrives at absolute concentrations by including *µ*_ox_(*T*).

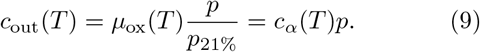

Here, *p*_21%_ is the reference value of oxygen partial pressure in the medium at an oxygen level of 21% in the gas phase. *c*_*α*_(*T*) is defined as a temperature-varying conversion factor.

| $T$ [°C] | 22 | 28 | 31 | 36 |
| --- | --- | --- | --- | --- |
| $c_{\alpha}(T)$ [$\mu\text{mol/l}$] | 1303.4 | 1166.3 | 1107.1 | 1019.3 |

### Subcellular ETC flux inference

The fluorescence decay curve is a histogram of photon counts *G* vs the photon arrival time *τ* obtained by recording the arrival time of each photon emitted from NADH autofluorescence. We model the normalized histogram *G*(*τ*) as the convolution of the normalized instrument response function (IRF) of the FLIM setup and the fluorescence decay of NADH with time, *F* (*τ*):

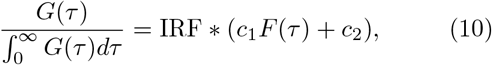

where *C*_1_ is the fractional contribution of the decay and *C*_2_ is the fraction of the background noise. The convolution is performed with periodic boundary condition to account for the sequential arrival of laser pulses with periodicity of 12.5ns. A time shift parameter is fitted to account for the shift between the peak of the IRF and the NADH decay curve. The IRF was measured as the normalized fluorescence decay curve of a urea crystal.

We assume a two-exponential decay model for NADH from the fact that enzyme-bound NADH displays a significantly longer lifetime than free NADH [15]:

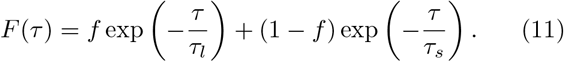

Here, *f* is the fraction of enzyme-bound NADH. The short fluorescence lifetime *τ*_*s*_ corresponds to free NADH, while the long fluorescence lifetime *τ*_*l*_ is associated with bound NADH.

To obtain the model parameters, *F* (*τ*) was convolved with the measured IRF to obtain the normalized *G*(*τ*) according to Eq.10. The normalized *G*(*τ*) was then fitted to the experimentally measured fluorescence decay curve after normalization to obtain *f, τ*_*l*_ and *τ*_*s*_. In addition, the fluorescence intensity *I* was obtained as the average number of photons collected within a single mitochondrial pixel per unit of scanning time.

We further assume that upon binding to the enzyme, the non-radiative decay rate of NADH decreases while the radiative decay rate stays constant. Since fluorescence lifetime is the inverse of the sum of the radiative and non-radiative decay rates, this indicates that the molecular brightness of NADH is proportional to its fluorescence lifetime, which enables the calculation of concentrations of bound NADH ([NADH_b_]) and free NADH ([NADH_f_]) from the fitted model parameters:

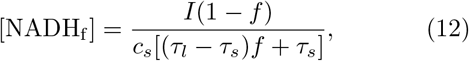

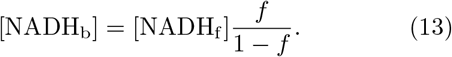

Here, *c*_*s*_ is an instrument-dependent calibration factor that relates fluorescence intensity to concentration. It was determined by titrating NADH and recording the FLIM signal in solutions. The validity of using Eqns 12 and 13 to measure free and bound NADH concentrations are established in Ref [19].

To infer NADH oxidation fluxes from free and bound NADH concentrations, we used the coarse-grained model of the NADH redox cycle inside mitochondria. The model predicts a relation between the total, steady-state flux through NADH oxidases in mitochondria as a function of concentration ratio of bound and free NADH (*β*) and free NADH concentration ([NADH_f_]):

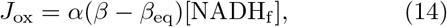

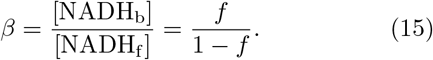

Here, *J*_ox_ is the steady-state NADH oxidation flux, which corresponds to the flux through the mitochondrial electron transport chain (ETC flux). *β* designates the ratio of bound to free NADH concentration, while *β*_eq_ is the corresponding value at zero flux conditions. Experimentally, *β*_eq_ was estimated as the lowest *β* during oxygen drop. The pre-factor *α* is related to the unbinding rates of NADH from the enzymes, and is assumed to be a constant. *α* was calibrated from direct measurement of oxygen consumption rate of the oocytes according to Eq.16.

FLIM collects fluorescence decay curves at each pixel with a pixel size of 420 nm, enabling the inference of NADH oxidative flux *J*_ox_ with subcellular resolution. To infer ETC flux as a function of distance to the oocytes center, FLIM curves from mitochondrial pixels in ten concentric rings of growing distance to the oocyte center were averaged together within each ring (Figure 1B). The double-exponential fitting was performed on the averaged NADH fluorescence decay curve within each ring to obtain *f* (*r*), *τ*_*l*_(*r*) and *τ*_*s*_(*r*) as a function of distance to the oocyte center. The NADH intensity *I*(*r*) was calculated within each ring by summing over all the photons from mitochondrial pixels within the ring and dividing by the number of mitochondrial pixels and scanning time. Free and bound NADH concentrations [NADH_f_](*r*) and [NADH_b_](*r*) were calculated within each ring according to equation 12 and 13. Finally, the spatially-dependent ETC flux *J*_ox_(*r*) was calculated according to equation 14, where *α* and *β*_*eq*_ are assumed to be spatially-independent.

### Calibration of ETC flux

To assign absolute value to the ETC flux, the prefactor *α* was calibrated by measuring the oxygen consumption rate (OCR) of the oocytes using a nanorespirometer (Unisense). In short, 10-15 oocytes were placed at the bottom of a glass caplillary well with a dimater of 0.68mm and a height of 3mm. After 1-2 hours of equilibration, a steady-state linear gradient of oxygen was formed in the well as a result of reaction-diffusion of oxygen. A motor-controlled Clark-type oxygen-sensitive electrode was used to measure the steady-state oxygen gradient in the well. The OCR of the oocytes was then caluclated according to Fick’s law OCR = *DA* ∇O_2_(*h*), where *A* is the cross-section area of the well and *D* is the diffusivity of oxygen. The whole respirometer is housed in a custom-built chamber for temperature control. OCR was measured at T=22°C, 28°C, 31°C and 36°C using the corresponding diffusivity of oxygen in water and solubility.

| $T$ [$^\circ\text{C}$] | 22 | 28 | 31 | 36 |
| --- | --- | --- | --- | --- |
| $\text{OCR}(T)$ [fmol/s] | 0.5475 | 0.7908 | 1.2110 | 2.3672 |

*α*(*T*) at different temperature *T* is calculated using the equation

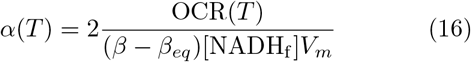

where *V*_*m*_ = 9.5 × 10^4^*µm*^3^ is the average volume of mitochondria per oocyte estimated from the MitoTracker measurement.

| $T$ [$^\circ\text{C}$] | 22 | 28 | 31 | 36 |
| --- | --- | --- | --- | --- |
| $\alpha(T)$ [1/s] | 1.1526 | 1.6648 | 2.5495 | 4.9836 |

### Inference of subcellular oxygen concentration

As discussed before, using the knowledge of subcellular oxidative flux *J*_ox_(*c*_out_, *r*) together with the assumption, that mitochondrial metabolism is the dominant sink for intracellularly diffusing oxygen, one can obtain subcellular oxygen profiles by integration according to eq. 3. This was done using custom code built around a standard finite-difference solver implemented in MatLab. As detailed in eq. 4, mixed boundary conditions were used.

### Fitting methods

The same Marquardt nonlinear least squares algorithm was used for oxidative flux inference from FLIM data fitting eq. (11) (Figure 1D-E and Figure 4B), estimation of decay length scales from *J*_ox_ and *ĉ* spatial profiles fitting modified Bessel functions described in sec. II B (Figure 2C and Figure 4C), fitting of Michaelis-Menten rate law to *J*_ox_(*ĉ*) (Figure 3A), estimation of approximate functional forms of Michaelis-Menten parameter profiles fitting linear and exponential functions (Figure 3B,C and Figure 4D, E).

## Code & Data availability

Together with the analysis pipeline used for creating the figures shown in the present work, the source code of the fitting algorithm is available on GitHub at https://github.com/MaxScw/mitoFluxGradients.

All data of the oxygen drop and temperature sweep experiment needed for running the code to recreate the figures can be downloaded from https://doi.org/10.25532/OPARA-1471.

## Acknowledgements

We thank Daniel Needleman for stimulating discussions. We thank Paul Stark for critical feedback on the manuscript. We thank Daniel Needleman and Yash Rana for proofreading the manuscript. This study received support from the Deutsche Forschungsgemeinschaft (DFG; German Research Foundation) under Germany’s Excellence Strategy — EXC-2068-390729961 — Cluster of Excellence Physics of Life of TU Dresden and from the French government under the France 2030 investment plan, as part of the Initiative d’Excellence d’Aix-Marseille Université - Amidex (AMX-23-CEI-064).

## Author contributions

X.Y. designed research; M.S. and X.Y. performed research; M.S., E.I., and X.Y. developed theoretical models; M.S. and X.Y. analyzed data; M.S., E.I., and X.Y. wrote the paper.

## Supplemental Material

### I. ROBUSTNESS OF OXYGEN PROFILES WITH RESPECT TO THE CHOICE OF OXYGEN DIFFUSION COEFFICIENT

In order to obtain subcellular oxygen profiles from integration of the oxidative flux *J*_ox_, the choice of a realistic diffusion coefficient across the cell is necessary. A natural choice is the diffusion of water at *T* = 36°C (*D* ≈ 3320 *µ*m^2^*/*s). To test the robustness of the inferred subcellular oxygen profile against the variations in the oxygen diffusivity, we varied the oxygen diffusivity within the range 2900 µm^2^*/s < D <* 3700 µm^2^*/*s and estimated the decay lengths of subcellular oxygen gradients at the lowest concentration of external oxygen level (*c*_out_ = 1.48*µ*M). The inferred decay length scale of the subcellular oxygen gradients lie in the range of 17.33 µm ≤ *λ*_ox_ ≤ 19.78 µm (Figure S1). It is therefore expected, that the results obtained based on the choice of *D* ≈ 3320 *µ*m^2^*/s* are robust.

**FIG. S1.**
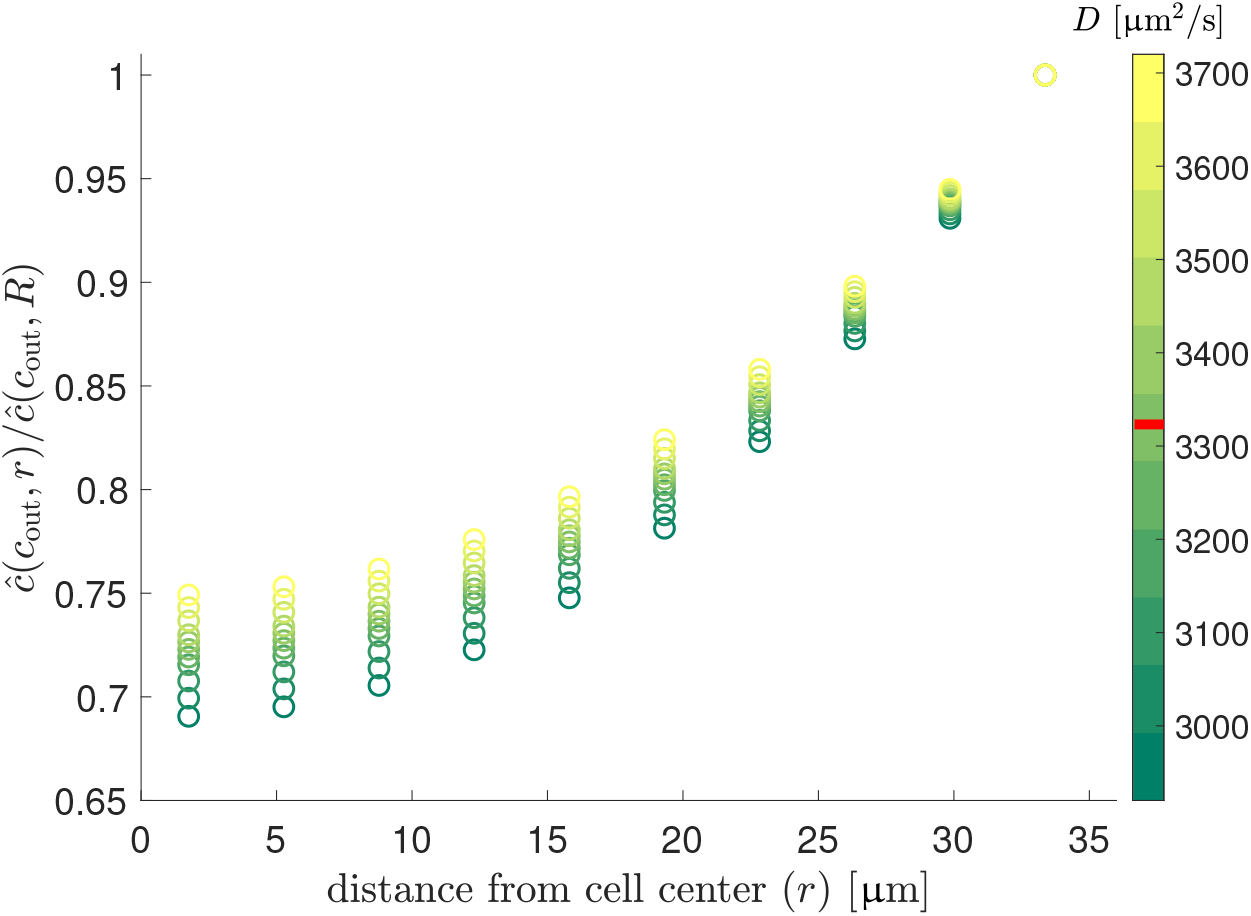
Subcellular oxygen profiles obtained from numerical integration of measured oxidative flux at different oxygen diffusivities (indicated in color, for calculation compare section II.B of the main text). The red line on the color scale indicates the diffusion coefficient that was used as a convention throughout all parts of the main text. All profiles are obtained from oxidative flux data at the lowest external oxygen concentration considered (*c*_out_ = 1.48 *µ*M). In the regime of low external oxygen, the strongest oxygen gradients across the cell are observed (diffusion-limited regime). Hence, the effect of varying oxygen diffusivity is expected to be most pronounced for the curves displayed.

### II. MODEL A: OXIDATIVE FLUX THROUGH OXYGEN REACTION-DIFFUSION WITH LINEAR KINETICS

From the observation, that the oxidative flux *J*_ox_ is proportional to the oxygen consumption flux within the cell, one can test a linear dependency of oxygen consumption flux on subcellular oxygen as a basic model. It turns out that this linear reaction-diffusion model (**model A**) is not sufficient to explain the observed variability of *J*_ox_ with external oxygen concentration (*c*_out_) without allowing for model parameters to depend on external oxygen concentrations.

We write the first-order reaction kinetics for oxygen consumption as

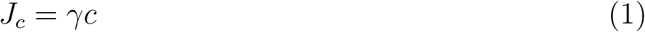

where *γ* is spatially uniform first-order reaction rate constant. At the steady state, one can write out the reaction-diffusion equation

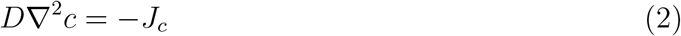

with boundary conditions

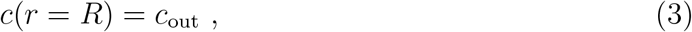

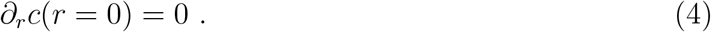

Simplifying the Laplacian due to spherical symmetry, one can obtain the analytical solution to Eqn. (2):

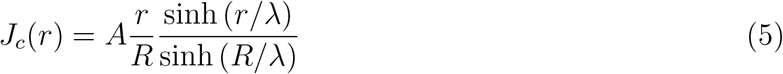

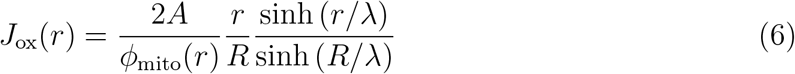

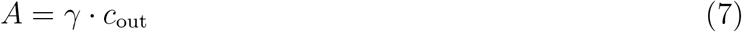

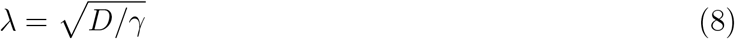

For *J*_ox_ gradients at different external oxygen levels, the functional form of Eqn.(6) was fit to the inferred ETC flux gradients (Figure S2A). The fitted parameters (*A* and *λ*) varied with external oxygen concentrations (Figure S2B and C), demonstrating the insufficiency of a linear reaction-diffusion model with constant parameters to explain the ETC flux gradients.

**FIG. S2.**
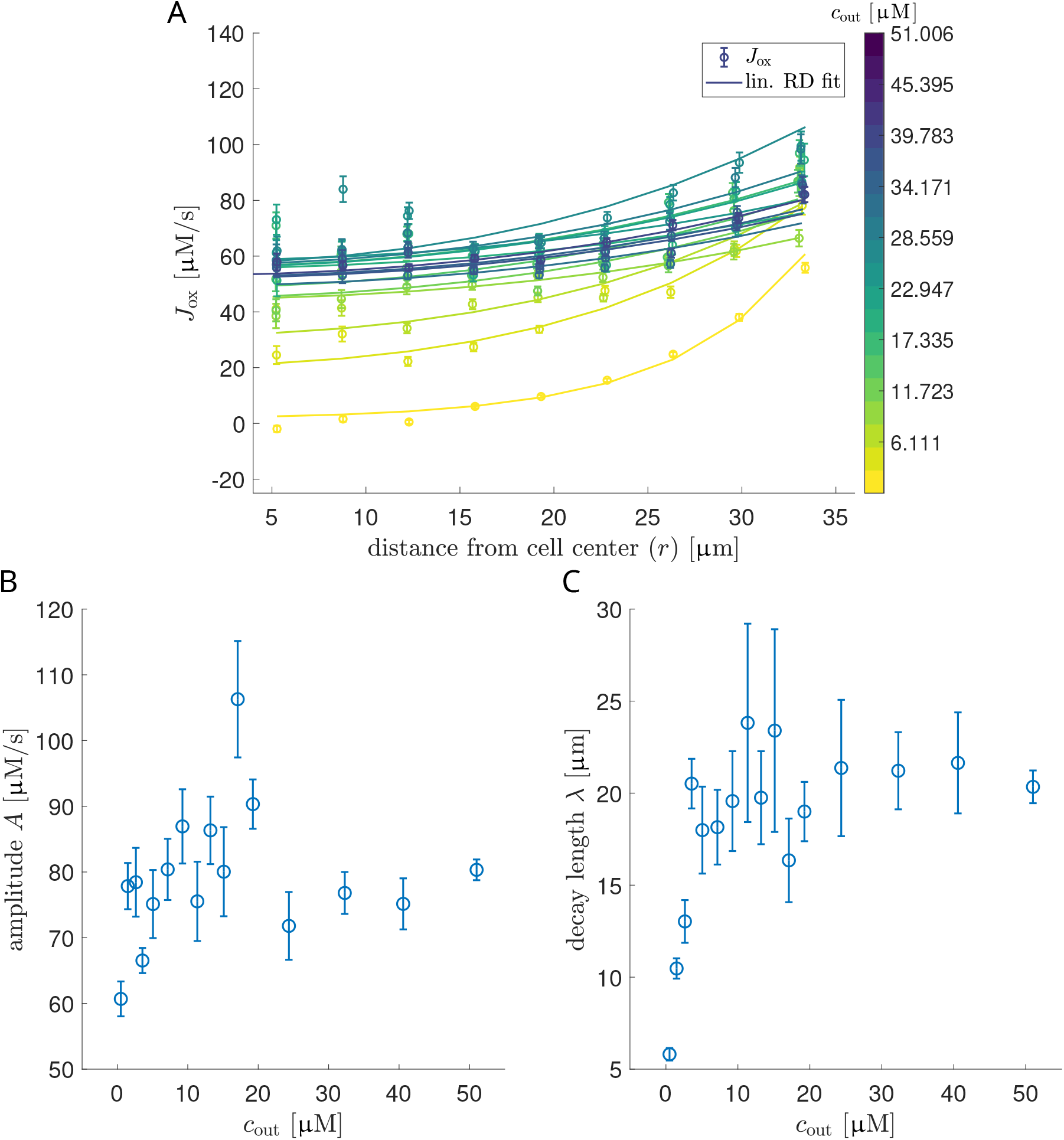
**A:** Fit of linear reaction-diffusion solution (Eqn. (6)) to inferred *J*_ox_ profiles. Error bars correspond to the standard deviation from the mean of *J*_ox_ over all oocytes. For each external oxygen concentration *c*_out_, the solution Eqn. (6) with unconstrained parameters *A, λ* is fit to the data. **B, C:** Optimal fit parameters of Eqn. (6) for all external oxygen concentrations considered. In order to fit the experimental data across all *c*_out_, both model parameters need to vary as a function of external oxygen concentration.

### III. MODEL B: OXIDATIVE FLUX THROUGH OXYGEN REACTION-DIFFUSION WITH SPATIALLY VARYING MICHAELIS-MENTEN KINETICS

As a linear dependency of the oxidative flux on subcellularly diffusing oxygen cannot explain the variability in *J*_ox_ without further assumptions, Michaelis-Menten like kinetics are arguably the simplest nonlinear form to test. It turns out, as indicated by the observation from Figure 2D in the main text that the subcellular ETC flux *J*_ox_(*ĉ*(*c*_out_, *r*), *r*) varies with *ĉ*(*c*_out_, *r*) in a spatially-dependent manner. Therefore, any nonlinear dependency of *J*_ox_ on oxygen alone cannot explain the experimental data completely. We thus need to account for the variability of *J*_ox_ across space by allowing the Michaelis-Menten model parameters to vary in space to fully explain the subcellular ETC flux gradients across all external oxygen levels. In this section, we explicitly derive the spatially-varying Michaelis-Menten model by allowing the binding (*k*_on_), unbinding(*k*_off_), enzyme catalysis rate (*k*_cat_) and the local enzyme concentrations ([*e*_tot_]) to vary in space (**model B**).

We consider a coarse-grained picture where mitochondria are considered as hosts of enzyme (*e*) that binds to oxygen (*c*) to form the complex (*e*^∗^*c*) before reducing oxygen to its respective product (*p*):

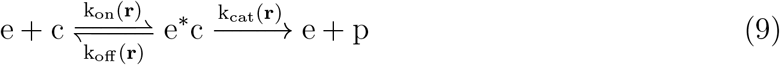

Here, [*e*] corresponds to the effective enzyme concentration (such as ETC complexes, while the known enzyme oxygen binds to is the Cytochrome c Oxidase, also known as the Complex IV), [*c*] is the concentration of subcellular oxygen and [*p*] is the reduced product from oxygen, such as H_2_O. *k*_on_ and *k*_off_ are the binding and unbinding rates of oxygen to the effective enzyme, while *k*_cat_ is the effective catalysis rate. Note that spatial variation of all kinetic parameters is allowed. However, we assume that enzymes are localized in mitochondria and neglect their transport. The reaction-diffusion equations of the metabolites and the enzymes are then given by:

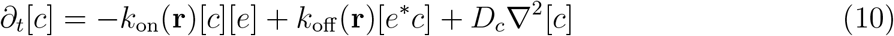

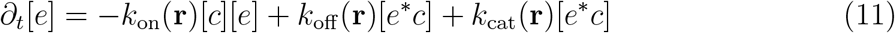

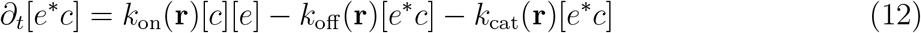

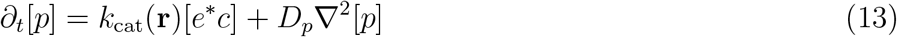

We define the local total concentration of the enzyme [*e*_tot_]

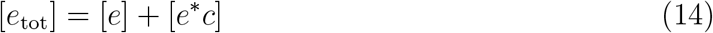

To account for the possible spatial heterogeneity of the mitochondrial composition, we allow [*e*_tot_] to vary in space. We then assume fast relaxation of substrate-enzyme complex concentration, i.e., *∂*[*e*^∗^*c*]*/∂t* = 0. This results using Eqs. (12),(14):

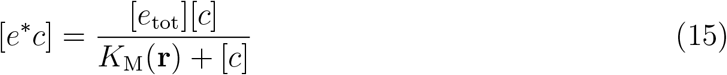

where the spatially varying Michaelis-Menten constant is defined as

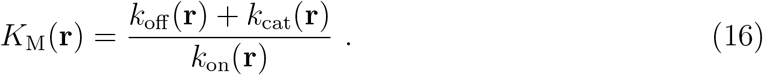

Inserting (15) into (10) and using (14) leads to

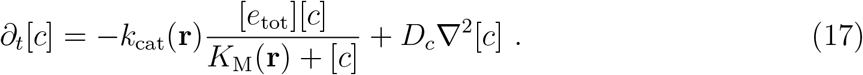

One thus arrives at a reaction-diffusion equation for oxygen, with a reaction term that is of Michaelis-Menten type and with spatially-dependent kinetic parameters. The maximal oxygen consumption rate can be identified as

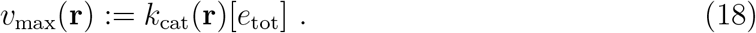

From the observation that NADH oxidation flux is proportional to the oxygen consumption rate of the mitochondria, we arrived at the theoretical prediction of *J*_ox_ as used in the main text:

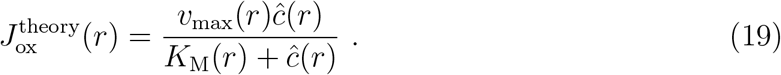

### IV. PROFILES OF FREE AND BOUND NADH CONCENTRATIONS AT DIFFERENT TEMPERATURES

**FIG. S3.**
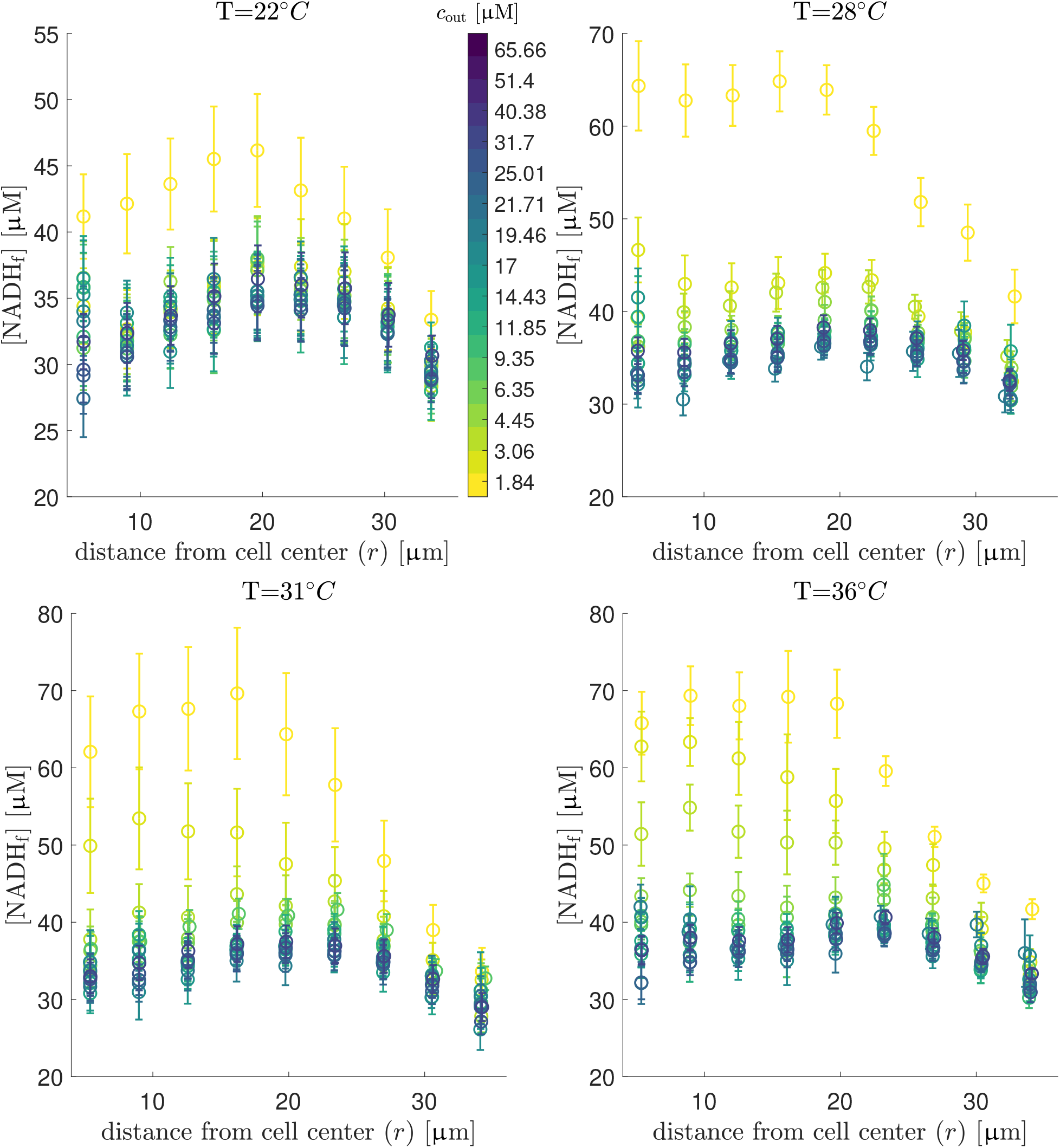
Spatial profiles of free NADH concentrations calculated based on spatially resolved FLIM measurement of NADH autofluorescence. Error bars correspond to the standard deviation of the mean obtained by averaging NADH profiles over all oocytes measured at the respective temperature.

**FIG. S4.**
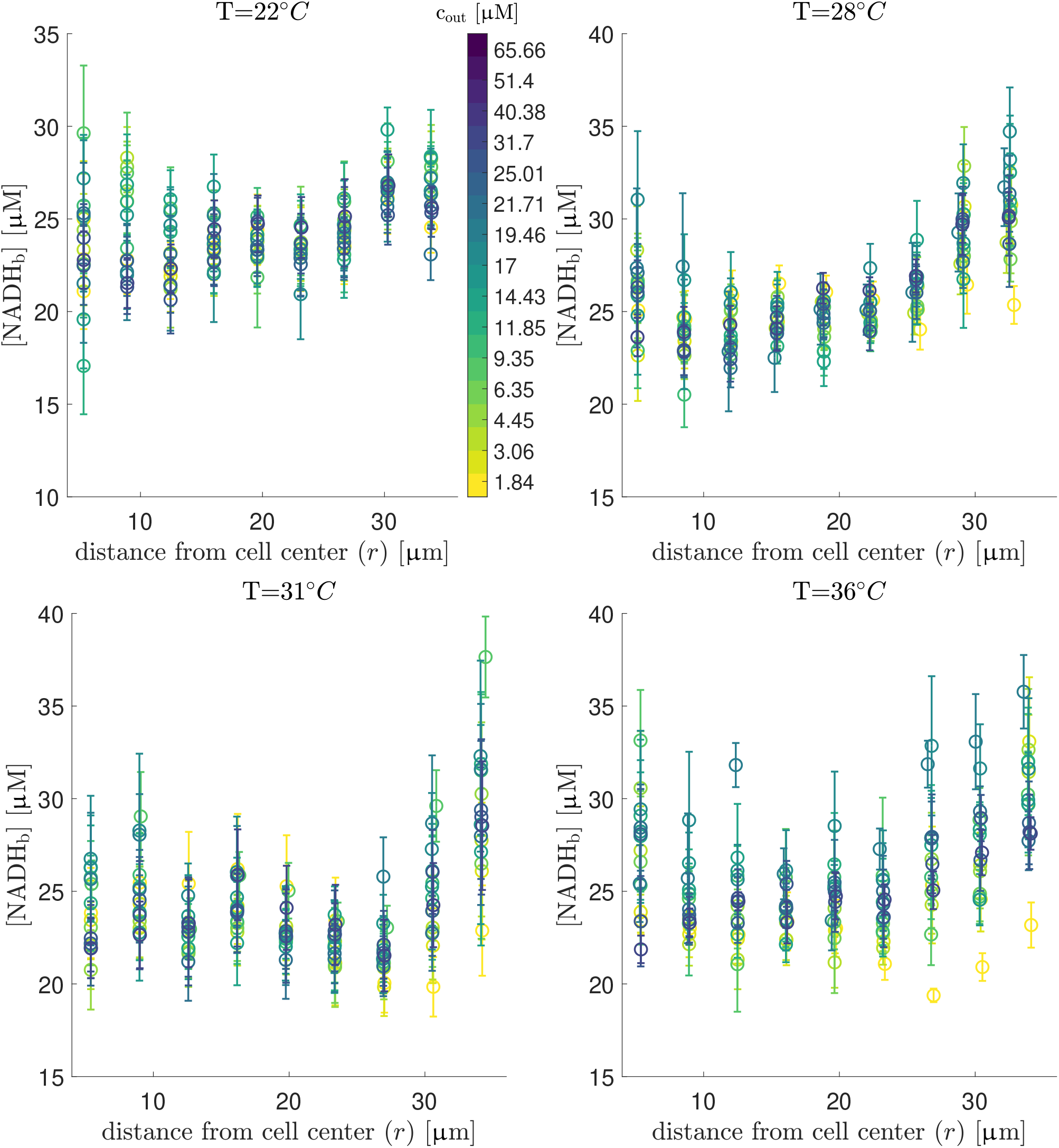
Spatial profiles of bound NADH concentrations calculated based on spatially resolved FLIM measurement of NADH autofluorescence. Error bars correspond to the standard deviation of the mean obtained by averaging NADH profiles over all oocytes measured at the respective temperature.

